# Coadapted genomes and selection on hybrids: Fisher’s geometric model explains a variety of empirical patterns

**DOI:** 10.1101/237925

**Authors:** Alexis Simon, Nicolas Bierne, John J. Welch

## Abstract

Natural selection plays a variety of roles in hybridization, speciation and admixture. Most research has focused on two extreme cases: crosses between closely-related inbred lines, where hybrids are fitter than their parents, or crosses between effectively isolated species, where hybrids suffer severe breakdown. Many natural populations must fall into intermediate regimes, with multiple types of gene interaction, but these are more difficult to study. Here, we develop a simple fitness landscape model, and show that it naturally interpolates between previous modeling approaches, involving mildly deleterious recessives, or discrete hybrid incompatibilities. The model yields several new predictions, which we test with genomic data from *Mytilus* mussels, and published data from plants (*Zea, Populus* and *Senecio*) and animals (*Mus, Teleogryllus* and *Drosophila*). The predictions are generally supported, and the model explains surprising empirical patterns that have been observed in both extreme regimes. Our approach enables novel and complementary uses of genome-wide datasets, which do not depend on identifying outlier loci, or “speciation genes” with anomalous effects. Given its simplicity and flexibility, and its predictive successes with a wide range of data, the approach should be readily extendable to other outstanding questions in the study of hybridization.

## 1 Introduction

Hybridization and admixture involve testing alleles in alternative genetic backgrounds. Most classical studies of hybridization can be placed into one of two classes. The first, involves crosses between closely-related inbred lines, where there is no coadaptation between the deleterious alleles that differentiate the lines, such that most hybrids are fitter than their parents. Wright’s single-locus theory of inbreeding was developed to interpret data of this kind (Crow 1952; Hallauer et al. 2010; Wright 1922, 1977). The second, involves crosses between effectively isolated species, where viable and fertile hybrids are very rare. Data of this kind are often analyzed by focusing on a small number of “speciation genes”, and interpreted using models of genetic incompatibilities (Coyne and Orr 1989; Dobzhansky 1937; Gavrilets 2004; Kalirad and Azevedo 2017; Orr 1995; Welch 2004).

The differences between these types of hybridization are clear, but it is equally clear that they are extremes of a continuum. Furthermore, the intermediate stages of this continuum are of particular interest, including, as they do, incipient speciation, and occasional introgression between partially-isolated populations (Duranton et al. 2017; Fraïsse et al. 2016a; Mendez et al. 2012; Waser 1993). However, it can be difficult to model natural selection in this intermediate regime, not least because models require a large number of parameters when they include epistatic effects between many loci. The empirical study of hybrid genotypes in this regime is also difficult. The analysis of lab crosses often focuses on segregation distortions of large effect, and pairwise incompatibilities (Abbott et al. 2013; Coyne and Orr 2004). This QTL-mapping framework can miss small effect mutations (Noor et al. 2001; Rockman 2012), which are difficult to identify individually, but whose cumulative effect can be substantial (Boyle et al. 2017).

One promising approach is to use Fisher’s geometric model, which assigns fitness values to genotypes using a model of optimizing selection on quantitative traits (Fisher 1930; G. Martin and Lenormand 2006; Orr 1998; Welch and Waxman 2003). The tools of quantitative genetics have often been used to study hybridization (e.g. Demuth and Wade 2005; Fitzpatrick 2008; Lynch 1991; Melchinger 1987), but Fisher’s model is fully additive at the level of phenotype, and the “traits” need not correspond in any simple way to standard quantitative traits (G. Martin 2014; Schiffman and Ralph 2017). Instead, the goal is to generate a rugged fitness landscape, which includes a wide variety of mutational effect sizes and epistatic interactions, with a minimum of free parameters (Barton 2017; Hwang et al. 2017; Orr 1998).

Here, we build on previous studies (Barton 2001; Chevin et al. 2014; Fraïsse et al. 2016b; Schiffman and Ralph 2017), and use Fisher’s geometric model to study hybridization. We develop a simple random-walk approximation, and show that it can naturally interpolate between previous modeling approaches, which are appropriate for the two extreme types of hybridization. We then show how the model can account for surprising empirical patterns that have been observed in both regimes (Moehring 2011; Moran et al. 2017; Wright 1977). Finally, we show that the model yields several novel predictions, and test these predictions with a wide range of new and existing data sets (Table 1).

**Table 1:**
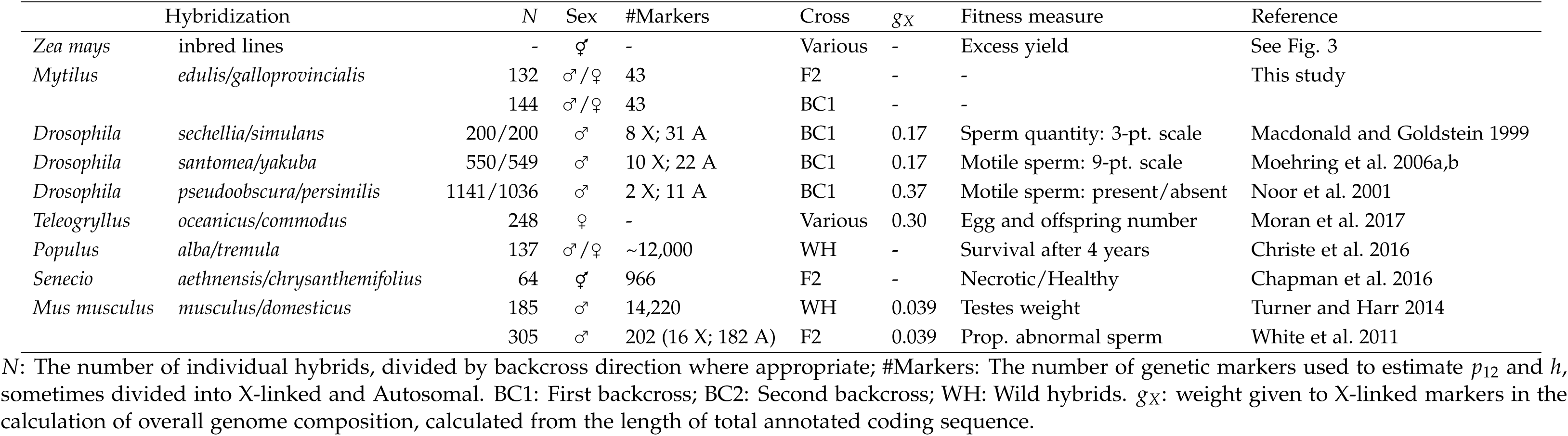
Data sets

## 2 Models and Results

### 2.1 The models

#### 2.1.1 Notation and basics

We will consider hybrids between two diploid populations, labeled P1 and P2, each of which is genetically uniform, but which differ from each other by *d* substitutions. The populations could generate 3^*d*^ distinct hybrid genotypes, but we are most interested in systematic differences between different types of hybrid (e.g., high versus low heterozygosity, males versus females, F1 versus F2 etc.). As such, following Turelli and Orr (2000), we describe hybrids using a “breakdown score”, *S*, which is larger for hybrids that are less fit. The relationship between *S* and fitness, *w*, might take a form such as

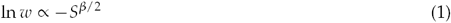

in which case, the parameter *β* adjusts the overall level of fitness dominance and epistasis, and can vary independently of other results (Fraïsse et al. 2016b; Hinze and Lamkey 2003; Tenaillon et al. 2007; see also Discussion). We now define the key quantity *f*, as the expected value of *S* for a particular class of hybrid, scaled by the expected value for the worst possible class.

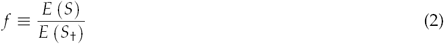

Here, *E* (*S*_†_), is the expected breakdown score for the class of hybrid with the lowest expected fitness. Therefore, *f* can vary between zero, for the best possible class of hybrid, and one, for the worst possible class.

To define classes of hybrid, we also follow Turelli and Orr (2000). We pay particular attention to inter-population heterozygosity, and define *p*_12_ as the proportion of the divergent sites that carry one allele from each of the parental types. We also define *p*_1_ and *p*_2_ as the proportion of divergent sites that carry only alleles originating from P1 or P2 respectively. Since *p*_1_ + *p*_2_ + *p*_12_= 1, it is convenient to introduce the hybrid index, *h*, which we define as the total proportion of divergent sites originating from P2.

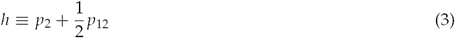

This quantity is closely related to measures of ancestry (e.g. Christe et al. 2016; Falush et al. 2003; Fitzpatrick 2012), although it considers only divergent sites. We can can now describe each individual genotype via its heterozygosity, *p*_12_, and hybrid index, *h*. Results below will mainly concern the dependency of *f* on *p*_12_ and *h*.

#### 2.1.2 Fisher’s geometric model

Fisher’s model is defined by *n* quantitative traits under optimizing selection (Fisher 1930). If the selection function is multivariate normal, including correlated selection, then we can rotate the axes and scale the trait values, to specify *n* new traits which are under independent selection of different strengths (G. Martin 2014; G. Martin and Lenormand 2006; Waxman and Welch 2005). An example with *n*= 2 is shown in Figure 1. We now define the breakdown score of a phenotype as

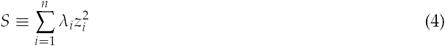

where, for trait *i, z*_*i*_ is its deviation from the optimum and *λ*_*i*_ is the strength of selection. By assumption, all mutational changes act additively on each trait, but their effects on breakdown can vary with the phenotype of the individual in which they appear, and this yields fitness epistasis. To specify the breakdown for each hybrid genotype, we would need to know the sizes and directions of all of the mutations that differentiate P1 and P2. However, a useful approximation is to treat the recombinant hybrid genotypes as if they lie along random walks in phenotypic space, where each fixed mutation contributes an expected *v*_*i*_ to the variance of the random walk on trait *i*. In this case, the worst possible class of hybrid will lie at the end of an unconstrained random walk away from the optimum, with no tendency for coadaptation among the changes. The walk can involve a maximum of *d* substitutions, and so we have

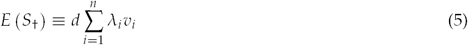

**Figure 1:**
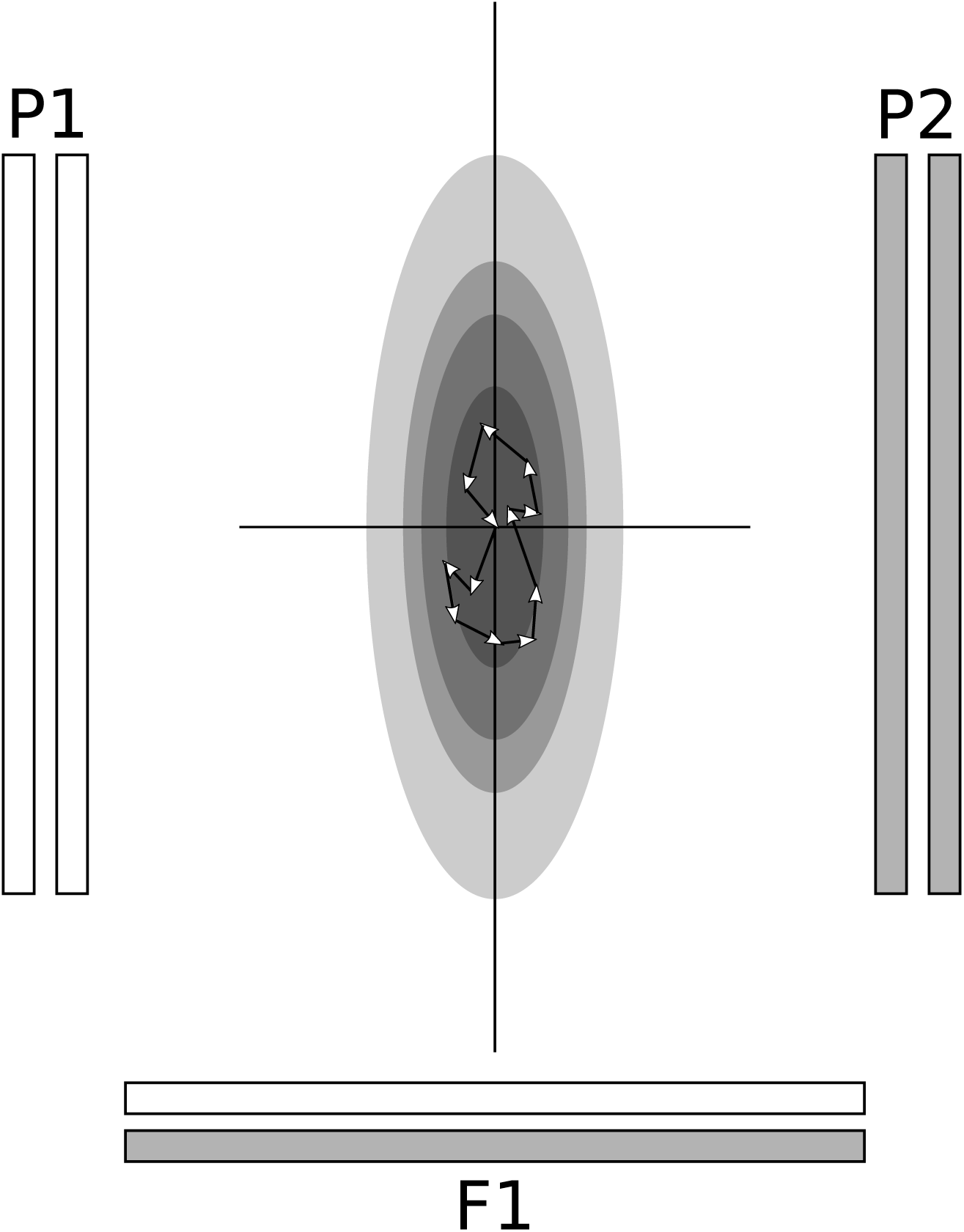
A schematic representation of Fisher’s geometric model, with *n*= 2 “traits”, each under optimizing selection of differing strengths. We consider hybrids between two diploid parental lines, P1 and P2, both of which have an optimal phenotype, but realized by different alleles. If we assume strict biparental inheritance, and additivity at the level of phenotype, then the initial F1 hybrid will have the midparental phenotype, which is also optimal. The expected fitness of other hybrids can be predicted by assuming that their component alleles form tethered random walks (Brownian bridges), between the three well-fit genotypes (see Appendix 1 for details).

Most hybrid genotypes will have higher fitness than this, because they contain combinations of alleles that are coadapted, as a result of past natural selection in their original backgrounds. To find the value of *f* (eq. 2) that applies to these genotypes, let us first assume that P1 and P2 are sufficiently well adapted, compared to the worst class of hybrid, to be treated as optimal. In this case, we fix *f* at zero for both parental types: *f*_P1_= *f*_P2_= 0. This implies that the midparental phenotype will also be optimal, and given the assumption of additivity, this midparent will be associated with the global heterozygote. We can now model the hybrid phenotypes as lying on a tethered random walk, or Brownian bridge, with these three optimal genotypes as fixed points; *f* is the variance associated with this Brownian bridge. In Appendix 1, we show that the result required is

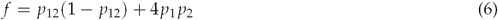

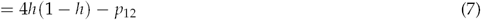

The fitness surface that is implied by eq. 7 is shown in Figure 2a. It should be noted that this prediction does not depend on any of the model parameters. For example, the number of traits, *n*, could affect the accuracy of the random-walk approximation (since *S* will tend to approach normality as *n* increases). But *n* does not appear in eq. 7, which depends on *p*_12_ and *h* alone.

**Figure 2:**
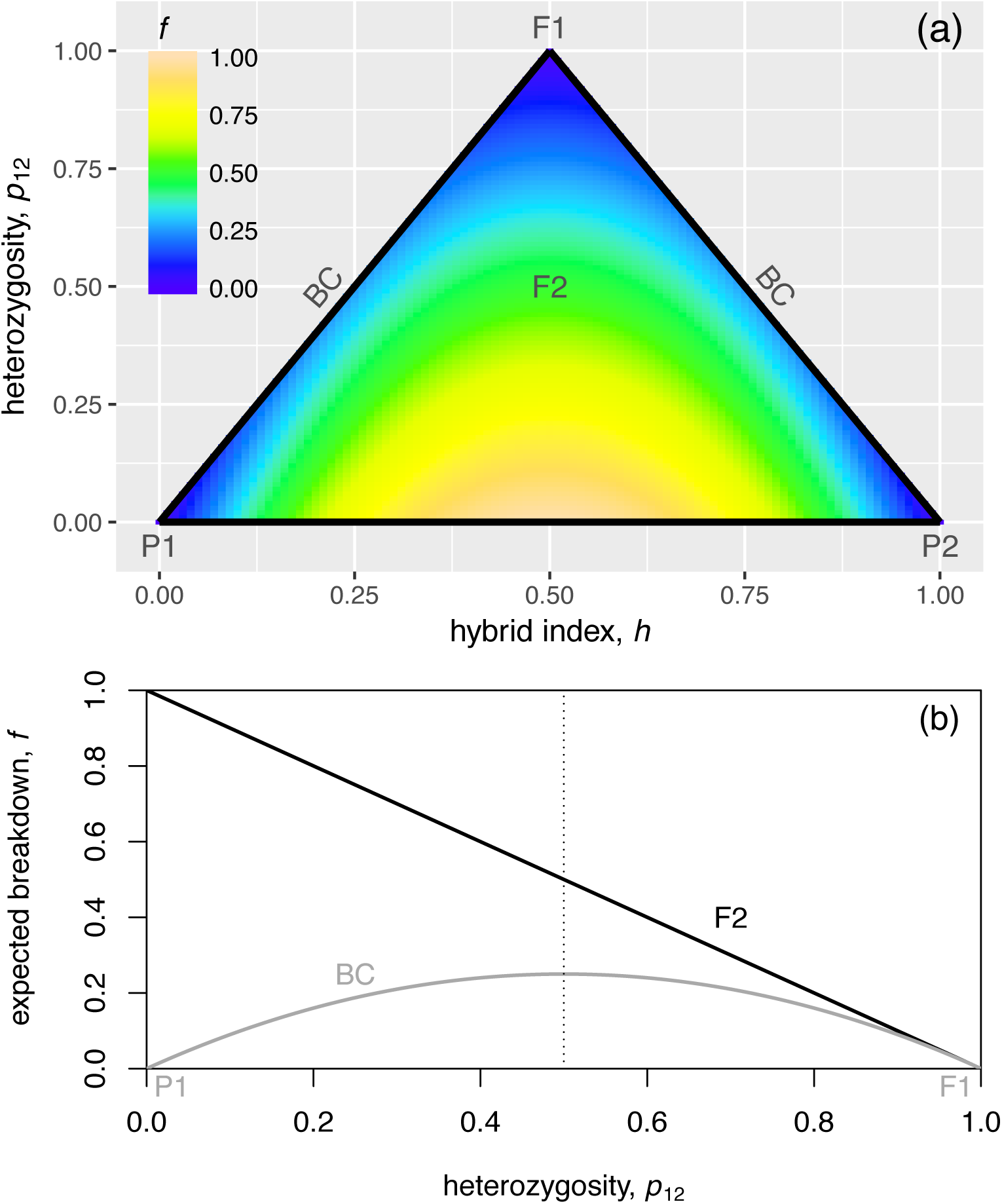
Panel (a) shows a heatplot of the fitness surface predicted by Fisher’s geometric model, for hybrid genotypes, when the parental types are well adapted (eq. 7 with *f*_P_= 0). The colors represent the relative expected breakdown score, *f*, with higher values corresponding to lower fitness (eqs. 1-2). Predictions are shown as a function of the interspecies heterozygosity, *p*_12_ and the hybrid index, *h* (eq. 3). The parental P1 and P2, are found at the lower corners, with *p*_12_= 0 and *h*= 0 or *h*= 1. With purely biparental inheritance, an initial F1 cross would be at the upper corner, with *p*_12_= 1, backcrosses would lie along the upright edges, with *h*= *p*_12_/2 or *h*= 1 *p*_12_/2, and the F2 would cluster in the center with 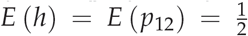. Panel (b) shows slices through this fitness surface, demonstrating that the selection on heterozygosity, *p*_12_, will differ according to cross type. For the F2 (F1 F1), heterozygosity is under directional selection, towards higher values. For backcrosses (such as BC1: F1 P1), then if the parental types are well adapted, heterozygosity is under symmetrical diversifying selection, away from the Mendelian expectation for the first backcross, and towards higher or lower values.

It is also possible to relax the assumption that the parents are optimally fit. In Appendix 1, we show that a simple and useful expression arises when the midparent is optimal, but the parents are suboptimal. This implies that both parents are equally maladapted: *f*_P1_= *f*_P2_ *≡ f* _P_, and we find that

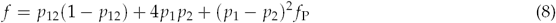

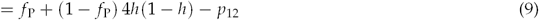

Fitness surfaces with varying levels of parental maladaptation are illustrated in Supplementary Figure S1. As expected, eq. 9 reduces to eq. 7 when *f*_P_= 0, while in the other extreme case, when *f*_P_= 1 we find,

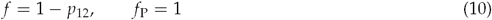

In this case, when parental fitness is no higher than the expected fitness of a random assembly of their alleles (eq. 2), then the breakdown is proportional to the total homozygosity (eq. 10). This result agrees with Wright’s (1922) single-locus theory of inbreeding, which was developed to analyze crosses between closely-related inbred lines (see also eq. 18 below). This agreement makes intuitive sense: when *f*_P_= 1, all of the divergence between P1 and P2 must comprise deleterious mutations with no coadaptation. In its general form (eq. 9), Fisher’s model shows how coadaptation between the parental alleles affects selection in the hybrids.

#### 2.1.3 A general model of incompatibilities

The previous section showed that Wright’s theory of inbreeding appears as a special case of Fisher’s geometric model, when *f*_P_= 1. In this section, we show that the other extreme case, with *f*_P_= 0, can also be derived via an alternative route, using a model of speciation genetics. We show that eq. 7 can be obtained from a model of genetic incompatibilities, each involving alleles at a small number of loci (Fraïsse et al. 2016b; Gavrilets 2004; Orr 1995; Turelli and Orr 2000; Welch 2004). The aim of this section is solely to compare the two modeling approaches. Empirical tests of eq. 9 follow in subsequent sections.

Following Orr (1995), let us assume that certain combinations of alleles, at *ℓ ≤ d* of the divergent loci, can be intrinsically incompatible, while all other combinations confer high fitness. By assumption, the pure species genotypes, and their ancestral states, must be fit, but all other combinations have a fixed probability *ε*_*ℓ*_ of being incompatible. Under this model, the expected breakdown score for the worst class of hybrid is proportional to the expected number of incompatibilities, and this was calculated by Welch (2004, eqs. 1-2):

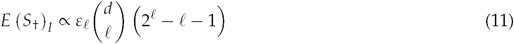

Here, and below, we use the subscript *I* to indicate a model of incompatibilities. To derive *f*_*I*_ (eq. 2), we note that hybrids will have higher fitness when some of the incompatibilities are absent from their genomes (Turelli and Orr 2000). The probability that an incompatibility is present depends on how many of the *ℓ* loci are heterozygous. For a genotype comprising *i* loci that are homozygous for the P1 allele, *j* loci homozygous for the P2 allele, and *k* loci that are heterozygous, the probability required is:

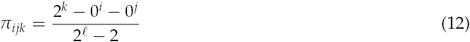

which is the proportion of the possible combinations of heterospecific alleles that are present in an “*ijk*” genotype. Incompatibilities may also have reduced effects due to recessivity, when their negative effects are masked by the presence of alternative, compatible alleles (Turelli and Orr 2000). To model this, we introduce the free parameter *s*_*ijk*_, which is the expected increase in breakdown when an incompatibility appears in an *ijk* genotype. Finally, in a hybrid genome characterized by *p*_1_, *p*_2_ and *p*_12_, the trinomial expansion of (*p*_1_ + *p*_2_ + *p*_12_)^*ℓ*^, tells us how many *ℓ*-locus genotypes of each type it is expected to contain. Putting these together, we have

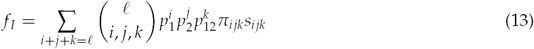

Equations 12-13 extend results with *ℓ*= 2 and *ℓ*= 3 from Turelli and Orr (2000), and represent a general model of breakdown caused by incompatibilities. A notable feature of these equations is their large number of free parameters. Even with symmetry between P1 and P2 (such that *s*_*ijk*_= *s*_*jik*_), we will still require a total of [*ℓ*(1 + *ℓ*/4)] different *s*_*ijk*_ values to specify the model (i.e., three extra parameters for two-locus incompatibilities, five parameters for three-locus incompatibilities etc.). There is good empirical evidence for, at least, two-three- and four-locus incompatibilities (Fraïsse et al. 2014), and so the full model would depend on at least 17 free parameters. By contrast, eq. 6, from Fisher’s geometric model, has no free parameters. The incompatibility-based model is therefore much more flexible, but also much more difficult to explore.

Because of this flexibility, however, it is also possible to find a set of *s*_*ijk*_ values that yield exactly the same dependencies as Fisher’s model. To do this, we set *f*_*I*_= *f*, using eqs. 6 and 13, and then solve for the *s*_*ijk*_. After some algebra, we find

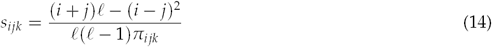

Equation 14 looks unwieldy, and it was derived solely to make the models agree. Nevertheless, in Appendix 2 and Supplementary Figure S2, we show that it embodies biologically plausible assumptions about incompatibilities, namely (i) partial recessivity, and (ii) increased levels of breakdown when incompatibilities are present with homozygous alleles from both parental lines. We further show that these *s*_*ijk*_ fall within the relatively narrow range of values that are required if the model is to yield a range of well-established empirical patterns (see also Turelli and Orr 2000). As such, when parental lines are well adapted compared to the worst possible class of hybrid (*f*_P_= 0), the key predictions of Fisher’s geometric model can also be derived from a general model of incompatibilities.

### 2.2 Testing the predictions with biparental inheritance

#### 2.2.1 Fitness differences between crosses

The simplest predictions from eq. 9 assume standard biparental inheritance at all loci. In this case, the standard cross types can be easily located on the fitness landscape shown in Figure 2a (Fitzpatrick 2012). With biparental inheritance, hybrids from the initial F1 cross (P1 *×* P2) will be heterozygous at all divergent loci 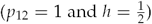; as such, eq. 9 predicts no breakdown for these F1.

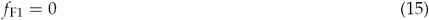

If the parental types are maladapted, then eq. 15 implies that *f*_P_ > *f*_F1_, and so there will be F1 hybrid vigor. Hybrid vigor can also appear at later crosses, but only when the parents are very maladapted. To see this, we can rearrange eq. 9 to provide a general condition for hybrid vigor:

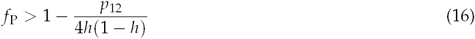

This condition will be much harder to satisfy for crosses beyond the F1. For example, in the first backcross (F1 *×* P1), all heterospecific alleles are heterozygous, and the expected heterozygosity is 50%: 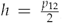, 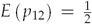, (Fitzpatrick 2012). As such, eq. 16 predicts hybrid vigor only when 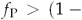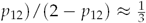. Conditions for hybrid vigor are even more stringent in the F2 (F1 *×* F1), when the expected hybrid index and heterozygosity are both 50%: 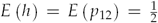. With these values, F2 hybrid vigor is predicted only when 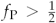. Taken together, these results predict that F1 vigor will be common, while hybrid breakdown will often appear in later crosses. This pattern has widespread empirical support (see references in Table A1 of Fraïsse et al. 2016b).

The model also makes quantitative predictions about the relative fitness of different crosses. Extensive data to test these predictions are available for *Zea mays*; these involve crosses of closely related and highly inbred lines, which do show hybrid vigor in the F2 and later crosses (Hallauer et al. 2010; Hinze and Lamkey 2003; Melchinger 1987; Neal 1935; Wright 1977). To analyze these data, a widely-used proxy for fitness is the excess yield of a cross, scaled by the excess yield of the F1. From eqs. 1-2, using a Taylor expansion the relevant quantity is approximately equal to

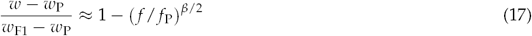

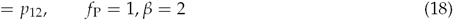

where *w* and *f* are the fitness and relative breakdown score for the hybrid of interest. For later crosses, these values will vary between individuals within a cross, due to segregation and recombination, but in this section we ignore this variation, and assume that *p*_12_ and *h* take their expected values for a given cross type. A fuller treatment is outlined in Appendix 3.

Equation 18 confirms that Fisher’s model reduces to Wright’s (1922) single-locus predictions for inbreeding, but only when all divergence is deleterious (*f*_P_= 1), and increases in breakdown score act independently on log fitness (*β*= 2). These single-locus predictions have strong support in *Zea mays* (Hallauer et al. 2010; Hinze and Lamkey 2003; Melchinger 1987; Neal 1935; Wright 1977). For example, as shown in Figure 3a, the excess yield of the F2 is often around 50%, equal to its expected heterozygosity (Hallauer et al. 2010; Wright 1977). It is also notable that the two outlying points (from Shehata and Dhawan 1975), are variety crosses, and not inbred lines in the strict sense.

**Figure 3:**
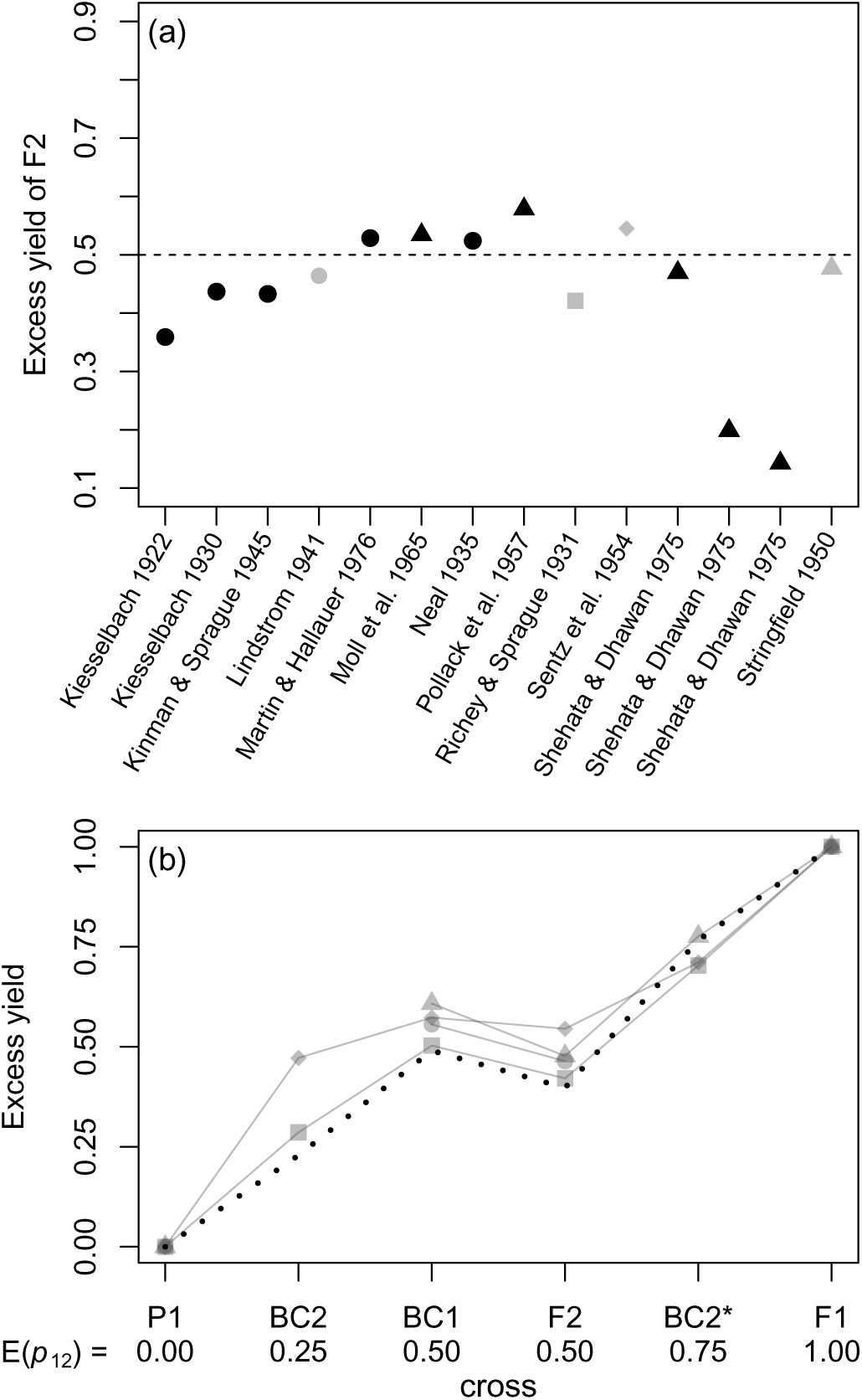
Data on hybrid vigor, from crosses of inbred *Zea mays*. The original data were collated by Wright (1977; see his Table 2.3.2), and Hallauer et al. (2010; see their Table 9.13), including only data from single crosses, where there was hybrid vigor in the F2, and yield was measured in quintals per hectare. Panel (a) plots the excess yield of the F2 (eq. 17). Results are shown for variety crosses (black triangles), as well as crosses of inbred lines in the strict sense (all other points). The dashed line shows the prediction of 0.5 from single-locus theory (eq. 18). Panel (b) shows the four data sets collated by Wright (1977), which allow us to compare the F2 and various backcrosses. These crosses, chosen to yield different levels of heterozygosity, are the parental type (P1), the second backcross (BC2= (F1× P1) × P1); the first backcross (BC1= F1 ×P1), the F2 (F1× F1), second backcross to the other parent (BC2= (F1× P1) × P2), and the F1 (P1 × P2) (The data of Stringfield 1950 replace BC2* with an F2 between two distinct F1, involving 3 distinct strains, but the predictions are unchanged). The grey symbols for the four data sets correspond to those used in panel (a). The dotted line in panel (b) shows predictions from Fisher’s model, assuming that the between-strain divergence contains limited coadaptation. The prediction uses eqs. 19-20 and 17, with *f*_P_= 0.75, and *β*= 2.5, which was chosen to fit the data of Richey and G. F. Sprague (1931). The model predicts both the roughly linear increase in vigor with heterozygosity, and the systematic difference between BC1 and F2.

Despite this predictive success, Wright (1977) noted a pattern that single-locus theory could not explain. In Wright’s words: “the most consistent deviation from expectation […] is the low yield of F2 in comparison with the first backcrosses” (Wright 1977, p. 39). Because 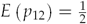 for both crosses, this difference must involve fitness epistasis. In fact, the pattern is predicted by Fisher’s model, when there is a small amount of coadaptation between the fixed alleles, such that 0.5 *< f*_P_ *<* 1 (see Supplementary Figure S1 for fitness surfaces of this type). To show this, Figure 3b plots the four relevant data sets collated by Wright, and compares the results to predictions from eq. 17 with *f*_P_= 0.75. The model predicts the roughly linear increase in vigor with mean heterozygosity, as with single locus theory, but also predicts the consistent difference in vigor between the backcross and F2.

**Table 2:** Tests for selection on heterozygosity in F2 and Backcrosses of *Mytilus* mussels.

|  |  |  |  |  |
| --- | --- | --- | --- | --- |
| Markers: | 43 | 43 | 33 | 33 |
| Data set: | All | No missing data | No missing data | Subsampled |
| Cross | $\hat{p}_{12}$ (N) $p$-value | $\hat{p}_{12}$ (N) $p$-value | $\hat{p}_{12}$ (N) $p$-value | $\hat{p}_{12}$ (N) $p$-value |
| F2 | 0.57 (132) $1.5 \times 10^{-6***}$ | 0.56 (88) $6.4 \times 10^{-4***}$ | 0.55 (91) 0.0033** | 0.56 (56) 0.0020** |
| BC | 0.57 (144) $1.3 \times 10^{-4***}$ | 0.53 (94) 0.0282* | 0.53 (105) 0.0569 | 0.52 (56) 0.5815 |

$\hat{p}_{12}$ : the estimated median heterozygosity; $N$ : the number of hybrid individuals sampled; $p$ -value: result of a Wilcoxon test of the null hypothesis median $p_{12} = 0.5$ . F2: random mating of F1 between *M. galloprovincialis* and *M. edulis*; BC: Backcross of the F1 to *M. galloprovincialis*. No missing data: all individuals with missing data for any of the markers were excluded; Subsampled: for the BC, any individual carrying a marker that was homozygous for the major allele carried by wild *M. edulis* populations was excluded; for the F2, we downsampled by sequencing order to equalize sample sizes.

**Table 3:**
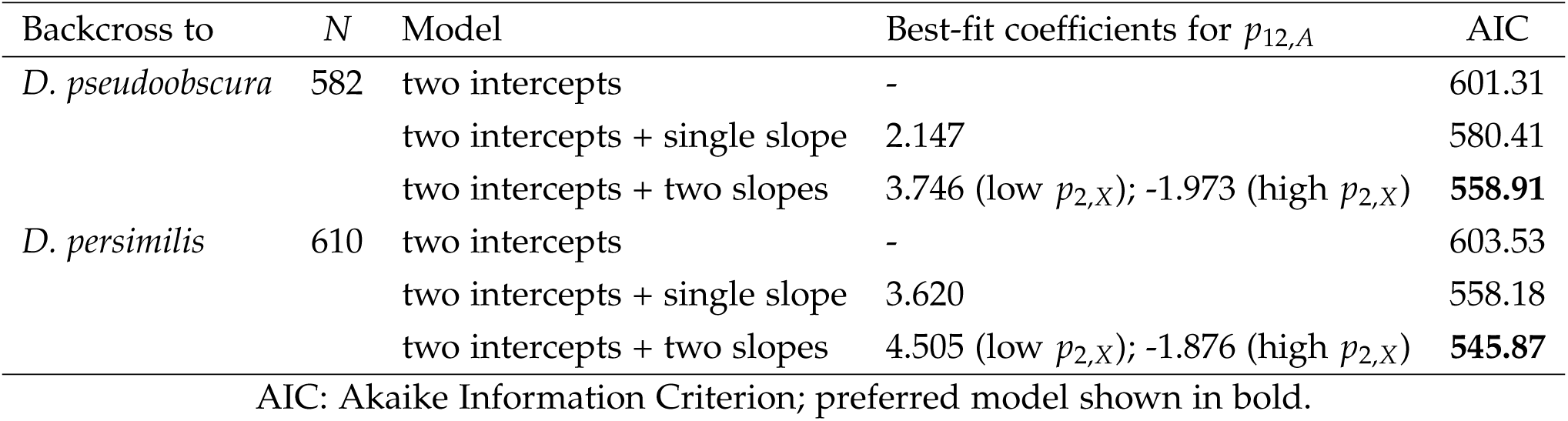
Regressions of male sterility on autosomal heterospecificity in *Drosophila* backcrosses

| Backcross to | $N$ | Model | Best-fit coefficients for $p_{12,A}$ | AIC |
| --- | --- | --- | --- | --- |
| *D. pseudoobscura* | 582 | two intercepts | - | 601.31 |
|  |  | two intercepts + single slope | 2.147 | 580.41 |
| | | two intercepts + two slopes | 3.746 (low $p_{2,X}$); -1.973 (high $p_{2,X}$) | **558.91** |
| *D. persimilis* | 610 | two intercepts | - | 603.53 |
|  |  | two intercepts + single slope | 3.620 | 558.18 |
| | | two intercepts + two slopes | 4.505 (low $p_{2,X}$); -1.876 (high $p_{2,X}$) | **545.87** |

AIC: Akaike Information Criterion; preferred model shown in bold.

#### 2.2.2 Selection on heterozygosity within crosses

In the previous section, we ignored between-individual variation in heterozygosity within a given cross type. In this section, we show how natural selection is predicted to act on this heterozygosity.

First, let us consider the F2. In this case, we have 4*h*(1 *− h*) *≈* 1 with relatively little variation between individuals (see Appendix 3 for details). Therefore, eq. 9 is well approximated by

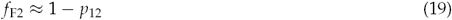

and so Wright’s result (eq. 10), applies in the F2, regardless of parental adaptedness. The prediction is that *p*_12_ will be under directional selection in the F2, favoring individuals with higher heterozygosity.

Now let us consider a backcross: F1 *×* P1. In this case, we have *p*_2_= 0, and so eq. 8 becomes

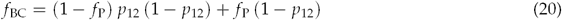

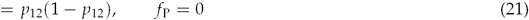

So selection in backcrosses varies with parental maladaptation. When *f* _P_ *>* 0.5 there is directional selection for higher heterozygosity, as in the F2. But when *f* _P_ is smaller, intermediate values of *p*_12_ yield the lowest expected fitness; when *f* _P_= 0, heterozygosity is under symmetrical disruptive selection, favoring heterozygosities that are either higher or lower than *p*_12_= 0.5 (eq. 21). These contrasting predictions are illustrated in Figure 2b (see also Supplementary Figure S1).

To test these predictions, we used a new set of genetic data from hybrids of the mussel species: *Mytilus edulis* and *Mytilus galloprovincialis* (Bierne et al. 2006, 2002). These species fall at the high end of the continuum of divergence during which introgression persists among incipient species (Roux et al. 2016). We used experimentally bred F2 and first backcrosses, with selection imposed implicitly, by the method of fertilization, and by our genotyping only individuals who survived to reproductive age (Bierne et al. 2006, 2002; see Methods and Supplementary Figure S3 for full details).

To estimate heterozygosity in each hybrid individual, we used the 43 markers that were heterozygous in all of the F1 hybrids used to make the subsequent crosses (see Supplementary Figure S3). We then asked whether the distribution of *p*_12_ values in recombinant hybrids was symmetrically distributed around its Mendelian expectation of *p*_12_= 0.5, or whether it was upwardly biased, as would be expected if directional selection were acting on heterozygosity. As shown in the first column of Table 2, Wilcoxon tests found that heterozygosities in surviving hybrids were significantly higher than expected, in both the F2 and backcross. These results may have been biased by the inclusion of individuals with missing data, because they showed higher heterozygosity (see Supplementary Table S1). We therefore repeated the test with these individuals excluded. As shown in the second column of Table 2, results were little changed, although the bias towards high heterozygosities was now weaker in the backcross.

Interpreting these results is complicated by the ongoing gene flow between *M. edulis* and *M. galloprovincialis* in nature (Bierne et al. 2002; Fraïsse et al. 2016a). To test for this, we genotyped 129 pure-species individuals, and repeated our analyses with a subset of 33 markers that were strongly differentiated between the pure species (see Methods, Supplementary Figure S3 and Supplementary Table S2 for details). With these markers, there was evidence of elevated heterozygosity in the F2, but not the backcross (Table 2 third column). We also noticed that many of our backcross hybrids, though backcrossed to *M. galloprovincialis*, carried homozygous alleles that were typical of *M. edulis*. We therefore repeated our analysis after excluding these “F2-like” backcrosses. Results, shown in the fourth column of Table 2, showed that the reduced BC data set showed no tendency for elevated heterozygosity. However, the bias towards higher heterozygosities remained in the F2, even when we subsampled to equalize the sample sizes.

Despite the problems of interpretation due to introgression and shared variants, the results support the prediction of eqs. 19-21: that directional selection on heterozygosity should act in the F2, but weakly or not at all in the backcross.

### 2.3 Predictions of Fisher’s geometric model with sex-specific inheritance

#### 2.3.1 Additional notation and basics

Results above assumed exclusively biparental inheritance. But the predictions of Fisher’s model are easily extended to include heteromorphic sex chromosomes, or loci with strictly uniparental inheritance, such as organelles or imprinted loci (Coyne and Orr 1989; Fraïsse et al. 2016b; Turelli and Moyle 2007; Turelli and Orr 2000). In these cases, *p*_12_, *p*_1_ and *p*_2_ are weighted sums of contributions from different types of locus. For example, with an X chromosome and autosomes, we have

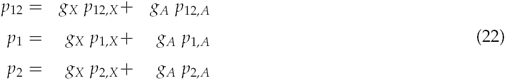

Here, the subscripts denote the chromosome type, so that *p*_12,*X*_ is the proportion of divergent sites on the X that are heterozygous, and *g*_*X*_ and *g*_*A*_ are weightings, which should sum to one (Turelli and Orr 2000). Results for specific cases can then be derived from eq. 8.

In the following sections, we apply this approach to three patterns involving sex-specific hybrid breakdown, in species with heteromorphic sex chromosomes. The first pattern, in the F1, is well known (Haldane 1922; Turelli and Orr 2000). The other patterns were observed in backcross data, which were generated to uncover the genetic basis of F1 breakdown, and test reasonable hypotheses about its causes (Moehring 2011; Moran et al. 2017). Both of these patterns have been called surprising, because neither agreed with the hypotheses (Moehring 2011; Moran et al. 2017). We show that all three patterns are consistent with predictions from Fisher’s geometric model.

#### 2.3.2 Haldane’s Rule

Haldane’s Rule states that sex-specific F1 breakdown usually appears in the heterogametic sex (Haldane 1922; Turelli and Orr 2000). To show how Fisher’s model predicts this pattern, we will assume an XO system for concreteness, such that females are homogametic, and males heterogametic. We will also assume that selection is identical in both sexes, and that pure-species males and females have the same fitness. These assumptions imply a form of dosage compensation, such that X-linked alleles have identical effects in homozygous or hemizygous state (Fraïsse et al. 2016b; Mank et al. 2011).

With these assumptions, the sole difference between male and female F1 is their heterozygosity. In XX females, all divergent sites are heterozygous, while in males, X-linked loci are hemizygous and from the maternal species, such that *p*_12_= 1 *− g*_*X*_ and *p*_1_ *p*_2_= 0. From eq. 8 we therefore find

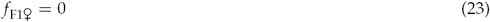

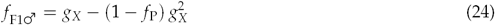

So *f*_F1♂_ > *f*_F1 ♀_ and Fisher’s model yields Haldane’s Rule (Barton 2001; Fraïsse et al. 2016b; Schiffman and Ralph 2017).

These results imply that female F1 will always have optimal fitness, regardless of the genetic distance between their parents (Barton 2001; Fraïsse et al. 2016b). However, if we extend the model, and allow for a proportion *g*_♀_ (*g* _♂_) of the divergence that is strictly maternally (paternally) inherited, then we find

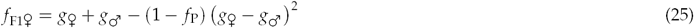

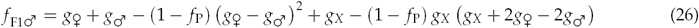

This still yields Haldane’s Rule for realistic parameter values (it always holds when *g*_*♀*_ *< g*_*A*_ for example), but breakdown can now appear in both sexes. This has two important consequences. First,exceptions to Haldane’s Rule can appear, but only in rare circumstances: when uniparentally-inherited loci act on traits that are subject to selection only in the homogametic sex (Fraïsse et al. 2016b). Second, from eqs. 2 and 5, the fitness of all F1 hybrids will tend to decline with genetic divergence; this yields an “F1 speciation clock” (Edmands 2002; Fraïsse et al. 2016b).

#### 2.3.3 Male backcrosses

A surprising pattern in backcross data was observed by Moehring (2011). Moehring reanalyzed three data sets of reciprocal backcrosses from *Drosophila* species, namely *D. simulans/sechellia* (Macdonald and Goldstein 1999); *D. santomea/yakuba* (Moehring et al. 2006a,b); and *D. pseudoobscura/persimilis* (Noor et al. 2001). In all three cases, F1♂ had low fertility, consistent with Haldane’s Rule, and so male hybrids were derived from the backcross F1♀ *×* P1♂ These backcross males vary in two measures of heterospecificity: their autosomal heterozygosity, *p*_12,*A*_, and their heterospecificity on the X, *p*_2,*X*_. Supplementary Figure S4 plots the data as a function of these two quantities.

Moehring (2011) predicted that sterility would increase with *p*_2,X_ and decrease with *p*_12,*A*_. These predictions follow from reasonable assumptions about Haldane’s Rule: that male sterility arises from partially recessive X-autosome interactions (Coyne and Orr 1989; Moehring 2011). Surprisingly, only one of these predictions was supported. In all six crosses, backcross sterility correlates strongly and positively with *p*_2,*X*_, but correlations with *p*_12,*A*_ are weak and inconsistent (see Supplementary Figure S4, and Table 3 of Moehring 2011).

Exactly this pattern is predicted by Fisher’s geometric model. To see this, Figure 4a depicts the fitness surface for hybrid males, as a function of *p*_2,*X*_ and *p*_12,*A*_. Individuals from a given cross might occupy a rectangular region, whose bounds are determined by *g*_*X*_. From annotated *Drosophila* genomes, we estimated that *g*_*X*_= 0.17 might characterize the *simulans/sechellia* and *yakuba/santomea* pairs, and that *g*_*X*_= 0.37 might characterize the *pseudoobscura/persimilis* pair (Table 1; see Methods for details). Figure 4 panels b-e show slices through the fitness surface for these values. In both cases, breakdown increases steadily with *p*_2,*X*_, except in the improbable case that the recombinant autosomes were completely heterozygous (Fig. 4b-c). This is consistent with the positive correlations observed. By contrast, the dependencies on *p*_12,*A*_ (Fig. 4d-e) vary in sign. This is consistent with the lack of consistent correlations with *p*_12,*A*_ (Supplementary Figure S4).

**Figure 4:**
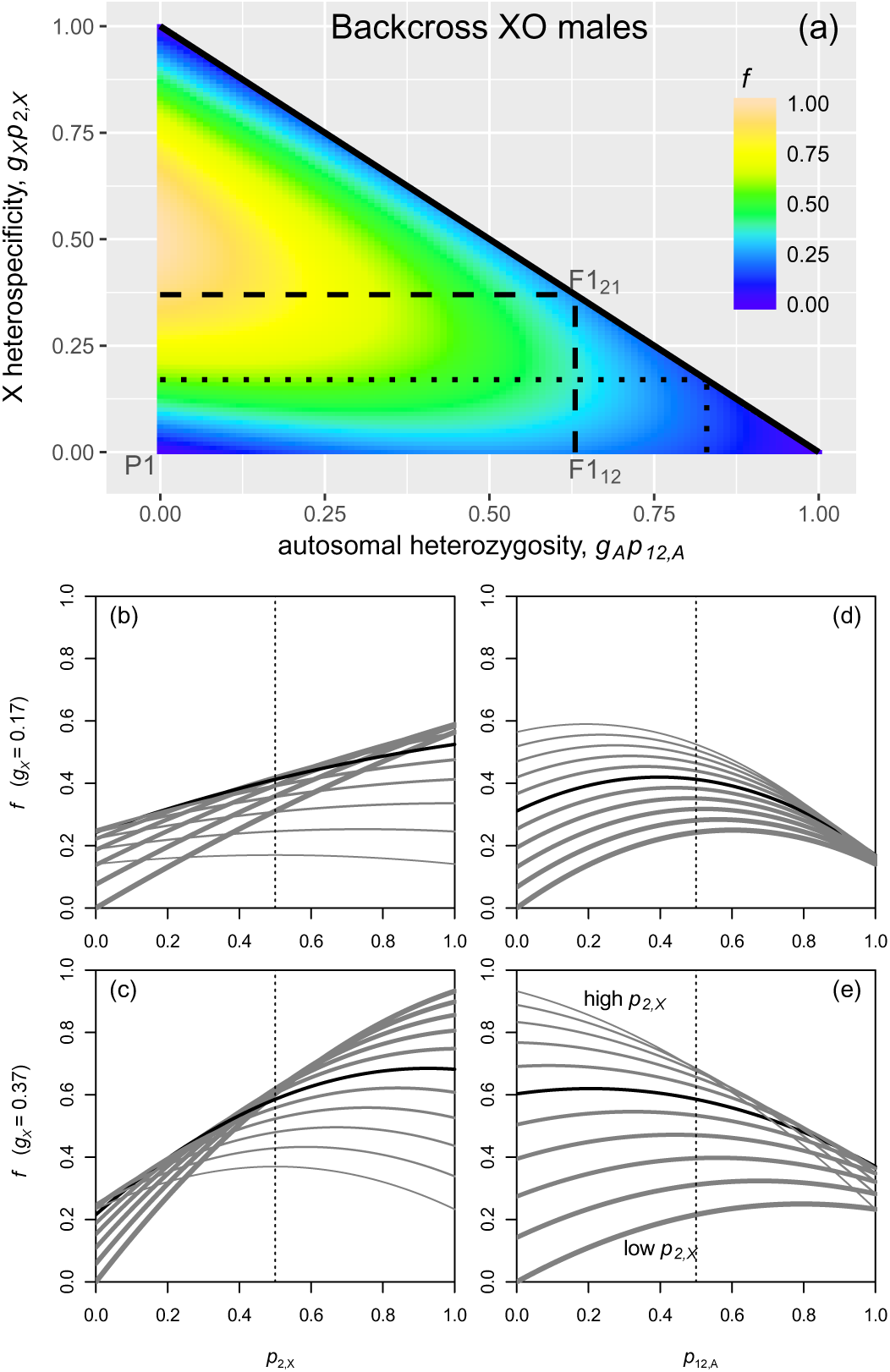
Predictions of Fisher’s geometric model for heterogametic male hybrids. For simplicity, the predictions neglect any contributions from uniparentally-inherited loci, and assume that the parental types are well adapted. For concreteness, we assume XO sex determination, so that hybrids differ in their autosomal heterozygosity, *p*_12,*A*_ and the proportion of alleles on the X that are heterospecific, *p*_2,*X*_. Panel (a) shows the fitness surface as a function of these two quantities. The dotted lines delimit the region that would apply to a species pair with *g*_*X*_= 0.17 (as we have estimated for *Drosophila simulans/sechellia* and *D. santomea/yakuba*), and the dashed lines delimit the region that would apply to a species pair with *g*_*X*_= 0.37 (as we have estimated for *Drosophila persimilis/pseudoobscura*). Panels (b)-(e) show slices through this fitness surface, with vertical dotted lines showing the Mendelian expectations for a first backcross. Panels (b) and (c) show the dependency on X-linked heterospecificity. Results are shown when the autosomal heterozygosity is equal to its Mendelian expectation of 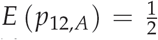 (black line), and over a range of values from *p*_12,*A*_= 0 (thickest gray line) to *p*_12,*A*_= 1 (thinnest gray line). Similarly, panels (d) and (e) show the dependency on autosomal heterozygosity, when the X-linked heterospecificity is at its expected value of 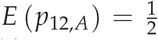 (black line), or over a range of values from *p*_2,*X*_= 0 (thickest gray line) to *p*_2,*X*_= 1 (thinnest gray line). Together, the plots show that Fisher’s geometric model can account for the surprising results of Moehring (2011), and generate a new supported prediction (Table 3).

Figure 4e also suggests a new testable prediction, which applies when *g*_*X*_ is large, as we have estimated for *D. persimilis/pseudoobscura* (Noor et al. 2001). When X-linked heterospecificity is low, then sterility is predicted to increase with *p*_12,*A*_, but when X-linked heterospecificity is high, then sterility is predicted to decrease with *p*_12,*A*_. With its two X-linked markers, the data of Noor et al. (2001) divide naturally into subsets with low, medium and high heterospecificity on the X (see the three rows of data points in Supplementary Figure S4e-f). Since we have no simple prediction when *p*_2,*X*_ is intermediate, we excluded these individuals, and then fit a GLM to the remaining data. We treated *p*_12,*A*_ as a linear predictor, and *p*_2,*X*_= 1 versus *p*_2,*X*_= 0 as a binary factor. In effect, we fit two linear regressions of sterility on *p*_12,*A*_, with different intercepts and slopes for the high-*p*_2,*X*_ and low-*p*_2,*X*_ individuals. As shown in Table 3, the predictions of Fisher’s model were supported for both backcross directions. Model selection favored a model with two slopes, and sterility correlated positively with *p*_12,*A*_ when *p*_2,*X*_= 0, and negatively with *p*_12,*A*_ when *p*_2,*X*_= 1.

#### 2.3.4 Female backcrosses

A second surprising pattern in backcross data was observed by Moran et al. (2017). These authors studied the field crickets *Teleogryllus oceanicus* and *T. commodus*, which have XO sex determination, and a large X chromosome (*g*_*X*_ *≈* 0.3; Moran et al. 2017). They are also a rare exception to Haldane’s Rule, with F1 sterility appearing solely in XX females (Hogan and Fontana 1973). Moran et al. (2017) hypothesized that female sterility might be caused by negative interactions between heterospecific alleles on the X, which appear together in F1_♀_, but not in F1♂.

To test this hypothesis, they generated backcross females with identical recombinant autosomes, but different levels of heterozygosity on the X, *p*_12,*X*_. The hypothesis of X-X incompatibilities predicts that fertility will decrease with *p*_12,*X*_, but this was not observed. The left-hand panels of Figure 5a reproduce the data of Moran et al. (2017), indicating crosses where X-heterozygosity was maximally high (*p*_12,*X*_= 1; circular points and darker shading), intermediate (*E* (*p*_12,*X*_)= 1/2; triangular points, and lighter shading), or maximally low (*p*_12,*X*_= 0; square points and no shading). The clearest trend is for backcrosses with intermediate levels of *p*_12,*X*_ to show the lowest fitness. By contrast, hybrids with extreme values of *p*_12,*X*_ showed no consistent differences.

**Figure 5:**
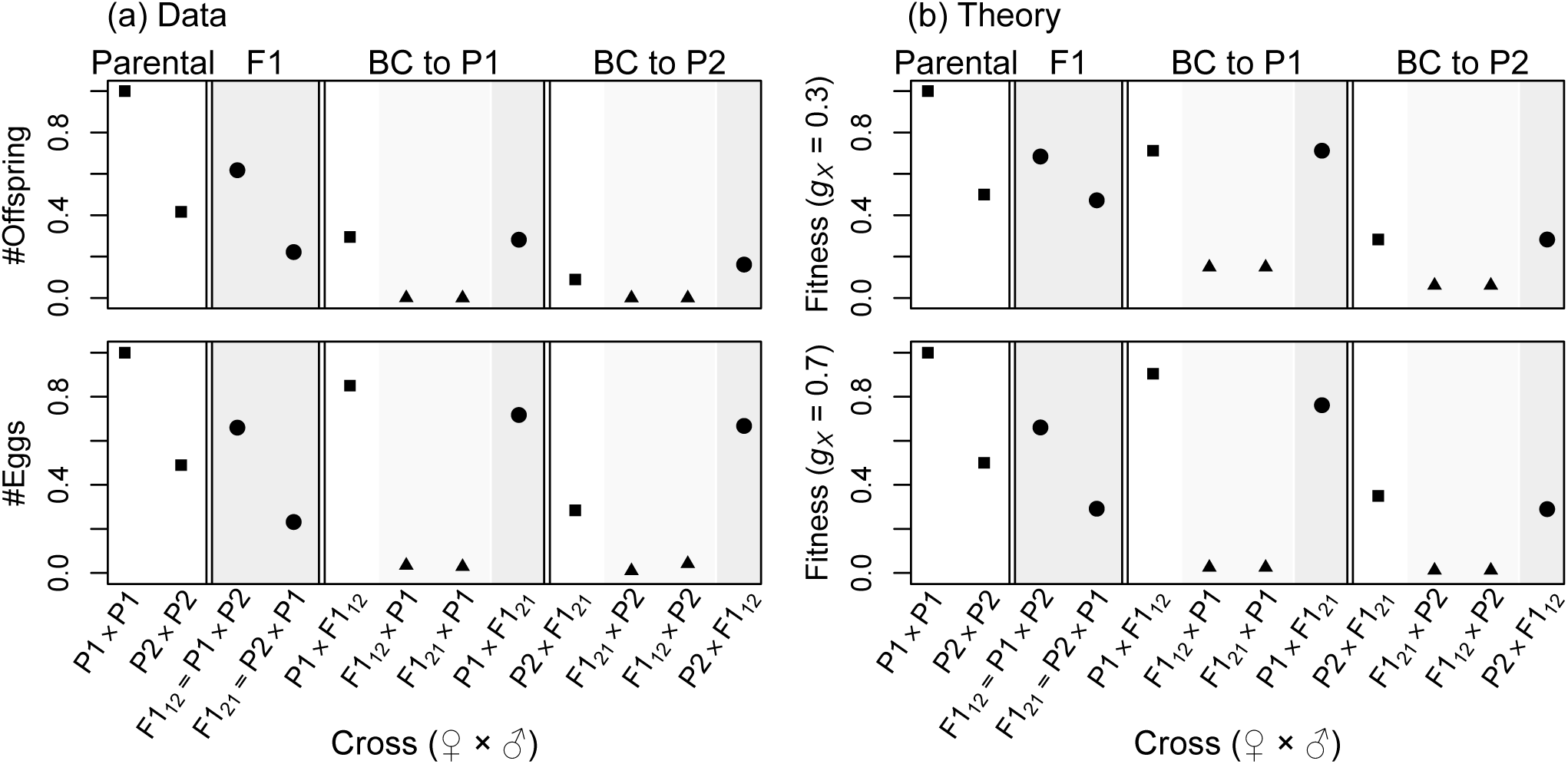
Data on female fertility, from crosses of the field crickets *Teleogryllus oceanicus* (P1), and *T. commodus* (P2), from Moran et al. (2017), compared to theoretical predictions from Fisher’s geometric model. Plotting styles denote the level of X-linked heterozygosity: high (circles and darker shading); intermediate (triangles and lighter shading) or low (squares and no shading). To plot the data (column (a)), we used the mean number of offspring per pair (upper panel), or mean number of eggs per pair (lower panel), each normalized by the value for P1 (see the supplementary information of Moran et al. 2017 for full details). The theoretical predictions (column (b)) are listed in Supplementary Table S3. In the upper panel, we assume *g*_*X*_= 0.3, as estimated from the chromosome sizes, and complete silencing of the paternal X (such that *π*= 1 in Supplementary Table S3). In the lower panel, we assume *g*_*X*_= 0.7, and incomplete silencing of the paternal X (*π*= 0.8), to improve fit to the egg data. While predictions apply to the rank order of fitnesses, to aid visualisation, we plot 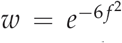 (see eq. 1), and set the parameter *f*_P2_ via 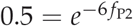, to reflect the lower fertility of this species under lab conditions.

To try and explain this pattern, let us note two clear asymmetries in the data. First, there are strong differences between the fitness levels of the two parental species in lab conditions, with *T. commodus* (labeled P2 in Figure 5) producing around half the eggs and offspring of *T. oceanicus* (P1 in Figure 5). Under Fisher’s geometric model, we can capture this asymmetry by assuming that P1 is optimal, while P2 is suboptimal. In Appendix 1, we show that these assumptions lead to the following result:

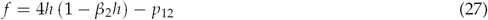

where

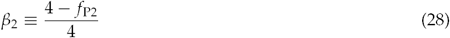

and *f*_P2_ is the breakdown associated with P2. Comparing eq. 27 to eq. 7 shows that maladaptation in one of the parental species introduces an asymmetry in the selection on the hybrid index, *h*, but leaves the form and strength of selection on heterozygosity *p*_12_ unchanged (see Supplementary Figure S1d).

Figure 5 shows a second asymmetry in the *Teleogryllus* data. The reciprocal F1 have very different fitness (Fraïsse et al. 2016b; Turelli and Moyle 2007). Because the X and autosomes will be identical for both cross directions, this asymmetry implies some sort of parent-of-origin effect on the phenotype. This could be either uniparental inheritance, or selective silencing (Fraïsse et al. 2016b; Turelli and Moyle 2007). One possibility is the speculation of Hoy and collaborators (see e.g., Butlin and Ritchie 1989; Hoy et al. 1977; Dr. Peter Moran pers. comm.), that dosage compensation in *Teleogryllus* involves silencing of the paternal X.

We can now use the foregoing assumptions to predict the levels of breakdown for each of the crosses produced by Moran et al. (2017). The predictions for each cross are listed in Supplementary Table S3, and plotted in the upper right-hand panels of Figure 5. This simple model explains several striking aspects of the *Teleogryllus* data.

Adjusting the parameter values can further improve the fit. For example, if we increase *g*_*X*_ (as would be case if divergent sites affecting female fecundity were clustered on the X), and assume that paternal X silencing is incomplete (affecting 80% of the divergent sites), then the results, shown in the lower-right-hand panel of Figure 5, agree well with the data on *Teleogryllus* egg number, as shown in the lower-left-hand panel (only the high fitness of P2 *×* F1_12_ is poorly predicted).

Further adjustments are possible, but these soon become *ad hoc*, at least without further knowledge of the true nature of parent-of-origin effects in *Teleogryllus*. The important point here is that Fisher’s geometrical model explains several key features of these hybrid data, while using only a single parameter derived from the data themselves; and even this parameter (*f*_P2_), was estimated from the parental control lines.

### 2.4 Estimating the fitness surface

Across a diverse collection of hybrids, equation 9 predicts that the hybrid index will be under symmetrical disruptive selection, and heterozygosity under directional selection. This prediction can be tested with data sets containing estimates of fitness, *h* and *p*_12_ for many hybrid individuals. Exactly such an analysis was presented by Christe et al. (2016), for families of wild hybrids from the forest trees, *Populus alba* and *P. tremula* (Christe et al. 2016; Lindtke et al. 2012, 2014). These authors scored survival over four years in a common-garden environment, and fit a GLM to these binary data (binary logistic regression, with “family” as a random effect), and predictors including linear and quadratic terms in *p*_12_ and *h*. Model selection favored a four-term model, with terms in *p*_12_, *h, h*^2^ (see Supplementary Table S5, and Supplementary information of Christe et al. 2016 for full details). For comparison with our theoretical predictions, we can write their best fit model in the following form:

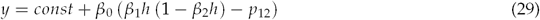

where *y* is the fitted value for hybrid breakdown. From eq. 9, Fisher’s model predicts that *β*_0_ *>* 0, 0 *≥ β*_1_ *≥* 4, and *β*_2_= 1, should hold. The best-fit model of Christe et al. (2016) corresponds to 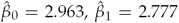 and 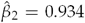, which supports the predictions of directional selection toward higher heterozygosity, and near-symmetrical diversifying selection on the hybrid index.

To obtain confidence intervals on these parameters, we fit the model of eq. 29 to the raw data of Christe et al. (2016). We also searched for other data sets, from which we could estimate the hybrid fitness surface. After applying some quality controls (see Methods, and Supplementary Table S1), we identified one other data set of wild hybrids, from the mouse subspecies *Mus musculus musculus/domesticus*, where male testes size was the proxy for fertility (Turner and Harr 2014). We also found four data sets of controlled crosses: F2 from the same mouse subspecies (White et al. 2011), and the ragworts *Senecio aethnensis* and *S. chrysanthemifolius* (Chapman et al. 2016); and the *Drosophila* backcrosses discussed above (Macdonald and Goldstein 1999; Moehring et al. 2006a,b). Unlike the data from wild hybrids, these single-cross data sets leave large regions of the fitness surface unsampled; nevertheless, they each contain enough variation in *h* and *p*_12_ for a meaningful estimation. Details of all six data sets are shown in Table 1, and they are plotted in Supplementary Figures S5-S7.

Figure 6 shows a summary of the estimated parameters, and full results are reported in Supplemen-tary Tables S4 and S5, and Supplementary Figures S5-S7. Taken together, the results show good support for the predictions of eq. 9. For all six data sets there was evidence of significant positive selection on heterozygosity (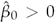 0 was preferred in all cases). Furthermore, for all six data sets, we inferred diversifying selection acting on the hybrid index. Estimates of *β*_2_, shown in the upper panel of Figure 6, show that this selection was near-symmetrical in all cases, such that 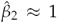. The poorest fit to the predictions was found for the *Drosophila* backcrosses, where estimates of *β*_1_ were significantly greater than the predicted upper bound of *β*_1_= 4 (Fig. 6 lower panel). But these data sets were least suited to our purpose, because estimates of *h* and *p*_12_ depend strongly on our rough estimate of *g*_*X*_= 0.17, and because they lack F2-like genotypes, from the center of the fitness surface (Figure 2a; Supplementary Figure S7). By contrast, results for the *Mus musculus* F2 (White et al. 2011), are remarkably close to the predictions of eq. 7 (Fig. 6; Supplementary Figure S6).

**Figure 6:**
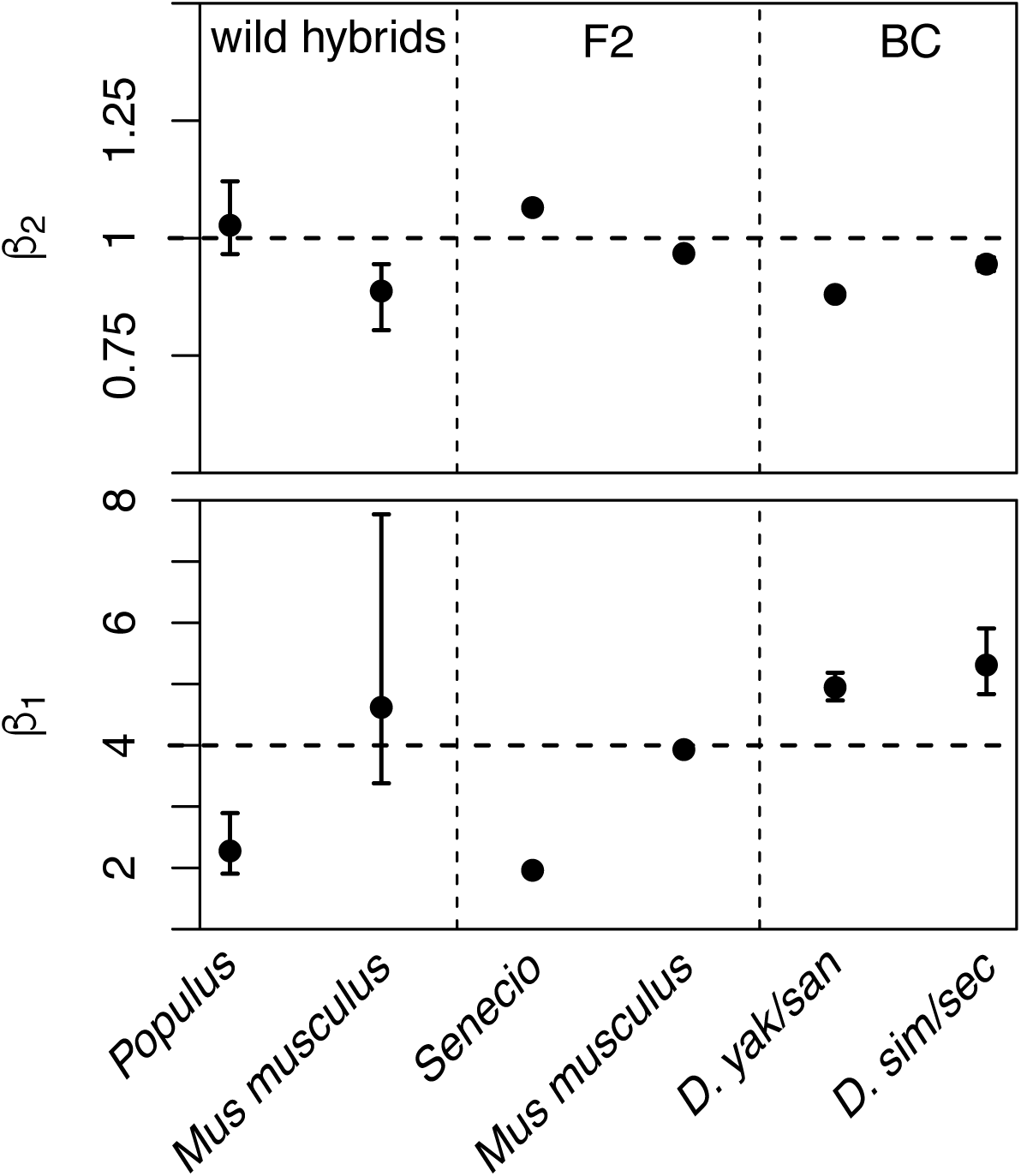
Best fit parameter estimates for the GLM of eq. 29, fit to fitness and genomic data from six data sets of hybrids (see Table 1 for details). The upper panel shows estimates of the coefficient *β*_2_ which determines the form of selection acting on the hybrid index, *h*. Estimates of *β*_2_= 1 are consistent with symmetrical diversifying selection. The lower panel show estimates of the coefficient *β*_1_ which determine the relative strength of selection acting on the hybrid index. Estimates of *β*_1_= 4 are predicted when the parental types are well adapted (eq. 7), while estimates 0 *< β*_1_ *<* 4 are predicted when the parental types are maladapted (eq. 9). Confidence intervals are defined as values that reduce the AIC by 2 units. These measures of uncertainty were not obtained for the F2 data, where variation in the hybrid index contributed little to the model fit, as predicted by eq. 19. Full details of the model fitting are found in the Methods and Supplementary Tables S4 and S5.

Two other features of the results deserve comment. First, for the two F2 data sets, it was not possible to provide meaningful confidence intervals for *β*_1_ and *β*_2_. This is because, for these two data sets, the terms in *h* and *h*^2^ did not make a significant contribution to model fit, and so the preferred model contained only selection on *p*_12_ (see Supplementary Table S5). This is consistent with our earlier prediction of eq. 19, and stems from the low variation in 4*h*(1 *− h*) among F2 hybrids (see Appendix 3 and Supplementary S5 and S6).

Second, for two of the data sets, *Populus* and *Senecio*, the estimates of *β*_1_ are substantially lower than 4 (Figure 6; Supplementary Figure S5). This is suggestive of parental maladaptation, creating true heterosis in the hybrids (see eq. 9). Consistent with this inference, there is independent evidence of parental load and F1 hybrid vigor in both species pairs (*Populus*: Caseys et al. 2015; Christe et al. 2017; *Senecio*: Abbott and Brennan 2014).

## 3 Discussion

In this article, we have used Fisher’s geometric model to develop predictions for the relative fitness of any class of hybrid. The modeling approach is simple, with few free parameters, and it generates a wide range of testable predictions. We have tested some of these predictions with new and published data sets (Table 1), and the major predictions of the model are well supported.

We emphasize that our approach is designed for coarse-grained patterns in the data, and typical outcomes of the evolutionary process, without considering the particular set of substitutions that differentiate the parental lines. The limitations of such an approach are seen in the low *r*^2^ values associated with our model fitting (Supplementary Table S4). Nevertheless, our approach should enable novel and complementary uses of genomic data sets, which do not depend on identifying individual loci with anomalous effects. Such a genome-wide interpretation of hybrid fitness is essentially lacking in the “speciation genes” framework.

A second goal of the present work was to show how Fisher’s model can interpolate between previous modeling approaches, namely the classical theory of inbreeding (Crow 1952; Wright 1922), and models of genetic incompatibilities, each involving a small number of loci (Dobzhansky 1937; Gavrilets 2004; Orr 1995; Welch 2004). We have also shown that Fisher’s model can account for empirical patterns that each approach has struggled to explain, although there are caveats to note in each case.

With inbred lines of *Zea mays*, we showed how observed differences in hybrid vigor are expected between the BC and F2 are expected, if we allow for a limited degree of coadaptation between the alleles that differentiate the lines (Figure 3; Wright 1977). The major caveat in this case is our simplifying assumption that the midparent is optimal (eq. 9; Appendix 1). This assumption is consistent with the *Zea* data, which show an enormous increase in F1 yield, but it is not clear how often the assumption would be met under an explicit model of mutation accumulation.

With the backcross data from *Drosophila* and *Teleogryllus*, the situation is more complicated. Moehring (2011) and Moran et al. (2017) showed that their data were not consistent with predictions from simple models of incompatibilities. But while these models were based on very reasonable assumptions, they only included incompatibilities of a single type (partially recessive X-A incompatibilities to explain Haldane’s Rule in *Drosophila*, or dominant X-X incompatibilities to explain the exception to Haldane’s Rule in *Teleogryllus*). We have shown that Fisher’s geometric model gives identical predictions to a general model of incompatibilities (eqs. 11-14), and that this general model can account for the patterns observed. It is also clear that the predictions were much more easily generated with eq. 6 than with eq. 13. In this case, there are two major caveats. First, the two models give identical predictions only when the dominance relations of incompatibilities are assigned in a particular way (eq. 14). But we have argued that these parameter values are biologically realistic, and strongly implied by other well-established empirical patterns (Appendix 2; Turelli and Orr 2000). Second, even when predictions are identical for the quantity *f* (eq. 2), the two approaches still make different predictions for other kinds of data, which were not considered in the present work. The most important difference is the dependency of log fitness on *d*, the genomic divergence between the species. Under Fisher’s geometric model, the log fitness of hybrids declines with *-d*^*β*^/^2^ (eqs. 1-2 and 5). By contrast, with the simplest models of incompatibilities, there is a snowball effect (Orr 1995), where the number of incompatibilities grows explosively with *d*^ℓ^ (eq. 11), and so log fitness declines with *-d*^*ℓβ/2*^. This is a genuine difference between the modeling approaches, although truly discriminatory tests may be difficult (Fraïsse et al. 2016b). For example, it may not always be possible to distinguish between a snowballing model with a low value of *β* (equivalent to strong positive epistasis between incompatibilities), or a model where *β* is higher, but where the number of “incompatibilities” does not snowball, because they appear and disappear as the genetic background changes (Fraïsse et al. 2016b; Guerrero et al. 2017; Kalirad and Azevedo 2017; Welch 2004).

Finally, given the simplicity and flexibility of the modeling approach explored here, and its predictive successes with a range of data, it should be readily extendable to address other outstanding questions in the study of hybridization. These include the putative role of hybridization in adaptive evolution (e.g. Duranton et al. 2017; Fraïsse et al. 2016a, b; Mendez et al. 2012), the effects of recombination in shaping patterns of divergence (Schumer et al. 2017), and the roles of intrinsic versus extrinsic isolation (Chevin et al. 2014). Given its ability to interpolate between models of different and extreme kinds, it should also be particularly useful for understanding hybridization in intermediate regimes, where parental genomes are characterized by both maladaptation and allelic coadaptation, or where the architecture of isolation involves many genes of small or moderate effect. Data — including those analyzed here — suggest that such architectures might be quite common (Baird 2017; Boyle et al. 2017; Buerkle 2017; Davis and Wu 1996; Maside and Naveira 1996; Morán and Fontdevila 2014).

## 4 Methods

### 4.1 Mytilus data

Conserved tissues from the mussel species, *Mytilus edulis* and *Mytilus galloprovincialis,* and their hybrids, were retained from the work of Bierne et al. (2006, 2002). As reported in those studies, *M. edulis* from the North of France were crossed with *M. galloprovincialis* from the French Mediterranean coast to produce F1 hybrids (five males and one female; Bierne et al. 2002). The F1 were then used to produce an F2, and sex-reciprocal backcrosses to *M. galloprovincialis* (which we denote here as BC_12_ and BC_21_; Bierne et al. 2006). In particular, oocytes from the F1 female were fertilized by the pooled sperm of the five F1 males producing F2 individuals, from which 132 individuals were sampled; oocytes from the F1 female were fertilized by pooled sperm of five *M. galloprovincialis* males to produce BC_12_, from which 72 individuals were sampled; and five *M. galloprovincialis* females were fertilized by pooled sperm from the five F1 males, producing BC_21_, from which 72 individuals were sampled. In addition to these hybrids, we also genotyped 129 individuals from “reference” populations of the two species, found in regions with relatively little contemporary introgression. In particular, we sampled *M. galloprovincialis* from Thau in the Mediterranean Sea; and sampled *M. edulis* from four locations in the North Sea and English Channel (The Netherlands, Saint-Jouin, Villerville and Réville). Full details of these reference populations are found in Supplementary Table S2.

In each case, gill tissues were conserved in ethanol at −20° C. DNA was extracted using a NucleaMag 96 Tissue kit (Macherey-Nagel) and a KingFisher™ Flex (ThermoFisher Scientific). We then genotyped all samples for 98 *Mytilus* markers that were designed from the data of Fraïsse et al. (2016a). The flanking sequences of the 98 SNPs are provided in Supplementary Table S6. Genotyping was sub-contracted to LGC-genomics and performed with the KASP™ chemistry (Kompetitive Allele Specific PCR, Semagn et al. 2014). Results are shown in Supplementary Figure S3. Many of the 98 markers are not diagnostic for *M. edulis* and *M. galloprovincialis*, and so we retained only the 43 that were scored as heterozygous in all 6 of the F1 hybrids. To obtain a reduced set of strongly diagnostic markers, we measured sample allele frequencies in our pure species *M. edulis* and *M. galloprovincialis* samples (pooling *M. edulis* individuals across the four sampling locations; Supplementary Table S2), and retained only markers for which the absolute difference in allele frequencies between species was >90%. This yielded the set of 33 markers used for the right-hand columns in Table 2. The “subsampled” data shown in the fourth column of Table 2, excluded any BC hybrid who carried the major allele typical of *M. edulis* in homozygous form. This yielded 56 BC hybrids. We then retained the first 56 F2 to be sequenced, to equalize the sample sizes.

### 4.2 Collation of published data

We searched the literature for published data sets combining measures of individual hybrid fitness, with genomic data that could be used to estimate *p*_1_, *p*_2_ and *p*_12_. In addition to those shown in Table 1, we also examined data sets that proved unsuitable for the sort of reanalysis presented here. These included data sets where the measure of fecundity or fertility took an extreme low value for one of the pure species, suggesting that it is not a good proxy for fitness (e.g. Orgogozo et al. 2006), data where the fitness proxy correlated strongly with a measure of genetic abnormality such as aneuploidy (Xu and He 2011), or data where the states of many markers could not be unambiguously assigned, for example, due to shared variation. Before estimating the fitness surface, we also excluded any data set where there was a highly significant rank correlation between the proportion of missing data in an individual, and either their heterozygosity, or fitness. For this reason, we did not proceed with reanalyses of the excellent data sets of Li et al. (2011), or Routtu et al. (2014) (see Supplementary Table S1 for full details). For our reanalysis of the *Teleogryllus* data of Moran et al. (2017), we did not consider data from the second backcross, to avoid complications due to selection on heterozygosity during the first backcross, and because of an anomalous hatching rate in the *T. oceanicus* controls. For our reanalysis of the *Mus musculus* F2 (White et al. 2011), we used a conservative subset of these data; we excluded any individual where any X-linked marker was scored as heterozygous (indicative of sequencing errors in heterogametic males; White et al. 2011), and controlled for variation in the uniparentally inherited markers, by retaining only individuals carrying *M. m. domesticus* mitochrondria, and the *M. m. musculus* Y. However, results were little changed when we used all 304 individuals with sterility data (Supplementary Table S5). Results were also unaffected when we used alternative proxies for fitness (Supplementary Table S5).

### 4.3 Estimation of *g*_*X*_ from annotated genomes

For taxa with XY sex determination (Table 1), the weightings *g*_*X*_ and *g*_*A*_, which determine the contribution of the X and autosomes to the overall constitution of the genome (eqs. 22), were estimated from the total length of coding sequences associated with each chromosome type, ignoring the small contributions from the Y and mitochondria. In each case, we obtained the longest protein product for each unique gene, and then summed their lengths, using a custom R script. The *g*_*X*_ values, shown in Table 1, were calculated as the total length of X-linked CDS divided by the total CDS length. For *Mus musculus*, we used the reference genome assembly “GRCm38.p5”. For *Drosophila simulans*, we used the assembly “GCA_000754195.3 ASM75419v2”, and for *Drosophila yakuba* “GCA_000005975.1 dyak_caf1”. For *Drosophila pseudoobscura*, the current annotation was downloaded from FlyBase release 3.04 Gramates et al. 2017). The .gtf file was then sorted and merged (combining overlapping coding sequences on each chromosome) using BEDTools (Quinlan and Hall 2010). Coding sequence lengths were calculated and summed over each chromosome, using custom awk commands.

### 4.4 GLM methods

The linear model results shown in Table 3, Figure 6, Supplementary Tables S4 and S5, and Supplementary Figures S5-S7, were all fit in R v. 3.3.2 (R Core Team 2016). For data sets with quantitative fitness measures (Turner and Harr 2014; White et al. 2011; Supplementary Figure S6) we used the standard general linear models, with Gaussian errors, and chose data transformations to reduce heteroscedasticity. For binary fitness data (Chapman et al. 2016; Christe et al. 2016; Noor et al. 2001; Table 3; Supplementary Figure S5), we used binomial regression with a logit link implemented in the *glm* function; and with ordinal fitness data (Macdonald and Goldstein 1999; Moehring et al. 2006b; Supplementary Figure S7) we used proportional odds logistic regression (Agresti 2003), implemented in the *polr* function. In these cases, the *p*-values shown in Supplementary Table S5 were calculated by comparing the *t*-value to the upper tail of normal distribution, as in a Wald test. For the non-Gaussian models, we also report McFadden’s pseudo-*r*^2^, defined as one minus the ratio of log likelihoods for the fitted and null models (McFadden 1974).

## Acknowledgments

We are very grateful to all of the authors of the original data reanalyzed here and especially grateful to the following, who provided clarifications or reformatted data: Luisa Bresadola, Prof. E. Charles Brummer, Dr. Mark A. Chapman, Dr. Camille Christe, Prof. Eric Hagg, Prof. Xionglei He, Prof. Christian Lexer, Prof. Xuehui Li, Prof. Amanda Moehring, Prof. Bret Payseur, Prof. Michael White, and Dr. Gavin Woodruff. We are also grateful to Dr. Andrea Betancourt for advice on processing the genome annotations, and to Dr. Peter Moran, who spotted a serious mistake in an earlier draft.

## Author contributions

AS and JW worked on the mathematical models and derivations. All authors designed the study, collected the data and wrote the manuscript.

## Appendix 1: the random walk approximation with suboptimal parental types

In this Appendix, we derive the random walk approximation for the breakdown score of a given hybrid genotype, under Fisher’s geometric model. Let us begin by describing the two parental phenotypes as *n*-dimensional vectors, denoted **z**_P1_ and **z**_P2_, which are equal but opposite deviations from the midparental phenotype, denoted as **z**_mp_. So if we define

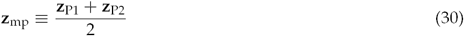

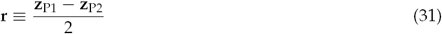

then

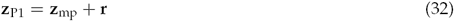

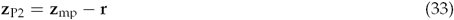

Below, we will use the notation *z*_mp,*i*_ and *r*_*i*_ to refer to the components of these vectors for trait *i*.

We can now consider the *d* mutations that differentiate P1 and P2 as describing equal but opposite paths from one of the parental phenotypes, to the midparent. Our approximation is to treat this path, on each of the *n* traits, as a Brownian bridge.

To derive this approximation, let *B*(*t*) denote a Brownian bridge, taking place over a single unit of time, and with a rate σ_*B*_. *B*(*t*) is normally distributed, with the following mean:

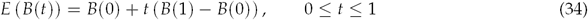

where B(0) and B(1) are the fixed points at the beginning and end of the random walk. The covariance at two time points is given by

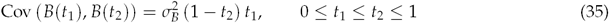

To model hybrid genotypes, we will need to count some sections of the random walk twice, to account for any homozygous alleles, and some sections only once, to account for any heterozygous alleles. Therefore, we are interested in the quantity:

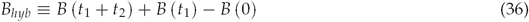

From eqs. 34-35, *B*_*hyb*_ will also be normal, with the following mean and variance

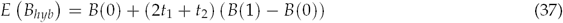

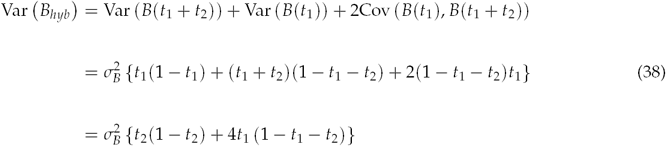

We can now apply these results to *z*_*i*_, the deviation from the optimum of trait *i* (see eq. 4 in the main text). In this case, the random walk begins from the trait value of parent P1: *B*(0)= *z*_mp,*i*_ + *r*_*i*_, ends at the midparent: *B*(1)= *z*_mp,*i*_, and has a rate equal to the total number of mutations, multiplied by their typical effect size: 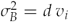. We then take the intermediate timepoints to be *t*_1_= *p*_2_ (this section of the walk is counted twice, to account for homozygous alleles), and *t*_2_= *p*_12_ (this section is counted once, to account for heterozygous alleles).

Putting these results together, we find that *z*_*i*_ is a normally distributed random variable, with the following properties:

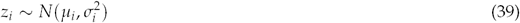

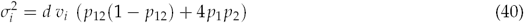

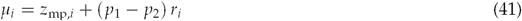

From eq. 4, the breakdown score, *S*, depends on the squared trait values, and from normal theory, we have

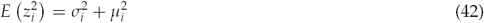

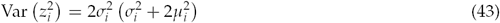

As such, *S* will be approximately gamma distributed, with a mean and variance given by the weighted sum of these quantities.

Let us now consider the special cases discussed in the main text. First, and simplest, is the case where both parents, and therefore the midparent, are phenotypically optimal. This implies that all *z*_mp,*i*_= 0 and all *r*_*i*_= 0, such that all *µ*_*i*_= 0. We then find

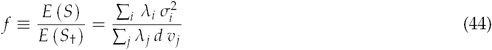

And this yields eqs. 6-7 of the main text. Next, let us consider the case where the midparent is optimal (all *z*_mp,*i*_= 0), but both parents are equally maladapted (some *r*_*i*_ *>* 0). In this case, 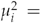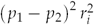 and 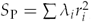 and so we find:

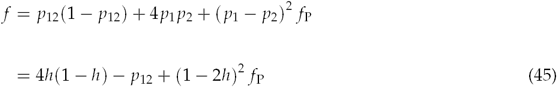

which yields eqs. 8-9 of the main text.

Let us finally consider the case that was used to analyse the data from *Teleogryllus*. In this case, we assume that one of the parental species (P2) is maladapted, while the other (P1) is optimal. Accordingly, we can set *z*_P1,*i*_= 0 and *z*_P2,*i*_= −2*r*_*i*_, such that *z*_mp,*i*_= *−r*_*i*_. We now have *µ*_*i*_= *−r*_*i*_ (1 + *p*_2_ *− p*_1_), and so

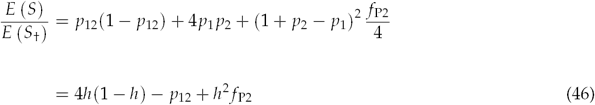

which yields eq. 27 of the main text. This situation is illustrated in Supplementary Figure S1d.

## Appendix 2: The dominance relations of incompatibilities

In this Appendix, we consider incompatibility-based models of hybrid fitness (eqs. 11-13). We examine different ways of assigning the parameters, *s*_*ijk*_, which appear in *f*_*I*_ (eq. 13), and represent the expected contribution to hybrid breakdown of individual incompatibilities, and especially, their dominance or recessivity (Turelli and Orr 2000). To understand this, let us begin by assigning the following functional form:

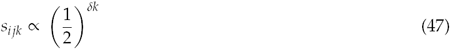

where below, we will use a constant of proportionality such that 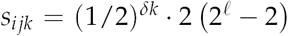 *to* simplify the results. In eq. 47, the parameter *δ* allows us to tune the dominance of incompatibilities, measured in terms of breakdown score, rather than fitness. When *δ*= 1, then each heterozygous locus halves the effects of incompatibility. This is equivalent to assuming that incompatibilities act multiplicatively, since each heterozygous locus halves the number of times that the incompatible combination of alleles is present in the genome. The *s*_*ijk*_ under multiplicative selection (*δ*= 1) are illustrated by the green points in Supplementary Figure S2.

To determine the predictions of this model, let us substitute eq. 47 into eq. 13, and set *δ*= 1. After some algebra, we find:

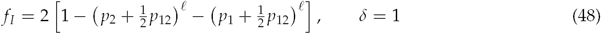

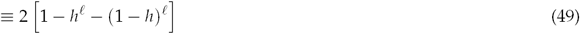

where *h* is the hybrid index, as defined in eq. 3. As such, when incompatibilities act multiplicatively, breakdown will depend solely on the total heterospecificity, and not at all on how the heterospecific alleles are arranged into genotypes (i.e. whether they appear as homozygotes or heterozygotes). It follows that breakdown is not predicted to change between the F1 and F2 crosses, and that homogametic F1, with 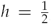, will have the highest possible breakdown score. As such, this multiplicative model cannot predict hybrid breakdown between the F1 and F2, or Haldane’s Rule, and both patterns are well supported (see Table A1 of Fraïsse et al. 2016b).

Now let us consider another extreme assumption. We assume that incompatibilities are fully recessive, such that no breakdown appears unless all incompatible alleles appear in homozygous or hemizygous form. We model this by making *δ* very large, such that *s*_*ijk*_= 0 unless *k*= 0. These values are illustrated by the red points in Supplementary Figure S2. With the assumption of complete recessivity, we find:

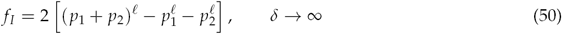

Equation 50 does not predict Haldane’s Rule, unless there is substantial uniparental inheritance from both the male and female parents. This is because *f*_*I*_= 0 if *p*_1_ *p*_2_= 0, and so both male and female F1 will have identical and optimal fitness. For similar reasons, eq. 50 predicts that the fitness of heterogametic backcrosses will decrease with *p*_12,*A*_: a prediction that is not supported by the relevant data (Moehring 2011).

We have shown that both extreme regimes (no recessivity, and complete recessivity) yield unsupported predictions. But what values of *δ* are biologically plausible? To answer this question, let us consider Haldane’s Rule under an incompatibility-based model, and ignoring uniparental inheritance. Assuming that males are heterogametic, Haldane’s Rule will hold when

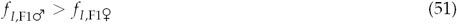

and using eqs. 13 and 47, after some algebra, eq. 51 is found to be equivalent to:

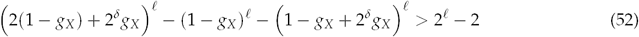

This condition is most difficult to satisfy when incompatibilities involve two loci (*ℓ*= 2), and in this case, we find the solution:

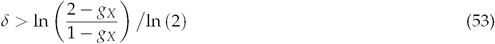

The value of *δ* that is required to yield Haldane’s Rule will therefore increase with *g*_*X*_. Towards the limit of the biologically plausible range, when two-thirds of the between-species divergence is X-linked (*g*_*X*_= 2/3) Haldane’s Rule will hold only if *δ >* 2. As such, setting *δ*= 2, such that each heterozygous locus reduces the breakdown score by a factor of four, will yield Haldane’s Rule in most cases. The *s*_*ijk*_ values from eq. 47 with *δ*= 2 are shown as yellow points in Supplementary Figure S2. Another feature of the model with *δ*= 2 is that it produces parameter dependencies that are very close to those predicted by Fisher’s geometrical model (see also Manna et al. 2011). The similarity is clearest with two-locus incompatibilities, where we find

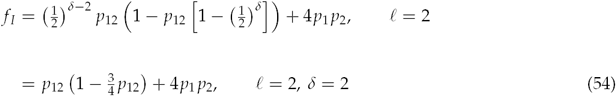

Comparing eq. 54 to eq. 6, shows that *f*_*I*_ *≈ f* when we use eq. 47 with *δ*= 2.

This is made even clearer when we compare the yellow points in Supplementary Figure S2, to the *s*_*ijk*_ values derived from eq. 14 of the main text, which were chosen to exactly match the predictions of Fisher’s geometric model (i.e. the values which yield *f*_*I*_= *f*). These *s*_*ijk*_ are shown as blue points in Supplementary Figure S2. The plot therefore clarifies the biologically-realistic assumptions embodied in eq. 14. First, these values reproduce the intermediate levels of recessivity that are required to generate Haldane’s Rule. Second, eq. 14 states that incompatibilities will have stronger effects when alleles from both parental species appear in homozygous state. For example, if the three alleles ABc form an incompatibility (with upper and lower case letters distinguishing alleles from P1 and P2), then eq. 14 predicts that the genotype Aa/BB/cc (with *ijk*= 111) will tend to have lower fitness than the genotype AA/BB/Cc (with *ijk*= 201) even though both genotypes contain the incompatibility, and both comprise two homozygous loci and one heterozygous locus.

## Appendix 3: Segregation and recombination

In this Appendix, we consider the effects of segregation and recombination on the expected levels of breakdown. For a recombinant cross, such as the F2 or backcross, *p*_12_ and *h* might vary between individuals, and so we must treat *f* as a random variable. To see this, let us write eq. 6 as

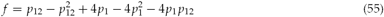

and so its expected value is:

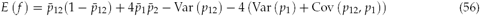

where overbars represent expected values. The variances and covariances in eq. 56 will depend on the distribution of the divergence across the genome, and on patterns of segregation and recombination. However, we can derive simple and useful predictions if we assume that the divergence is equally distributed among *m* freely recombining regions. In this case, we can use results from a multinomial distribution.

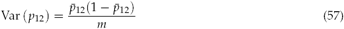

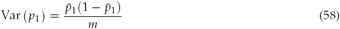

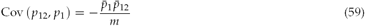

These results also apply to estimators from *m* independently segregating markers. As an example, let us compare the first backcross and the F2, in a population with strictly biparental inheritance. For the first backcross, we have 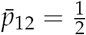 and *p*_1_ *p*_2_ = 0, and so

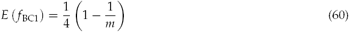

For the F2, we have 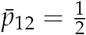 and 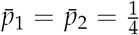, and so

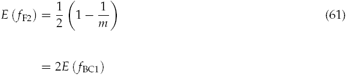

and so the predicted breakdown in the F2 is always double that of the first backcross. By the same method, we can also calculate the variance of *f*, but this requires higher order moments of the multinomial distribution (Newcomer et al. 2008), and this can yield lengthy expressions. However, the following results for the F2 are required to justify the approximation of eq. 19 from the main text.

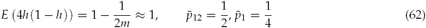

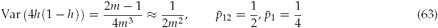

Comparing eq. 63 to eq. 57 shows that most of the variance in F2 breakdown will be due to variation in heterozygosity.

